# The Particle Filter Method to Integrate High-Speed Atomic Force Microscopy Measurement with Biomolecular Simulations

**DOI:** 10.1101/2020.03.12.988485

**Authors:** Sotaro Fuchigami, Toru Niina, Shoji Takada

**Affiliations:** Department of Biophysics, Graduate School of Science, Kyoto University, Kyoto 606-8502, Japan

## Abstract

The high-speed atomic force microscopy (HS-AFM) can observe structural dynamics of biomolecules at single-molecule level in real time near physiological condition, but its spatiotemporal resolution is limited. Complementarily, molecular dynamics (MD) simulations have higher spatiotemporal resolutions albeit with some artifact. Here, in order to integrate the HS-AFM data and coarse-grained (CG)-MD simulations, we develop a particle filter method, one of the sequential Bayesian data assimilation approaches. We tested the method in a twin experiment. We first made a reference HS-AFM movie from a CG-MD trajectory of a test molecule, a nucleosome, which serves as an “experimental measurement”. Then, we performed the particle filter simulation with 512 particles that captured large-scale nucleosome structural dynamics compatible with the AFM movie. Comparing the particle filter simulations with 8 - 8192 particles, we found that the use of more particles consistently results in larger likelihood for the whole AFM movie. By comparing the likelihoods from different ionic concentrations and from different timescales, we found that the “true” concentration and timescale can be inferred as the largest likelihood of the whole AFM movie, but not that of each AFM image. The particle filter method provides a general approach to integrate the HS-AFM data with MD simulations.

## INTRODUCTION

Biomolecular functions are, in general, realized via their structural dynamics. Albeit their central importance, characterizing protein structures both at high spatial and temporal resolutions remains one of the major unsolved challenges. On the one hand, X-ray crystallography, cryo-electron microscopy, and others can give structure models at high spatial resolution, i.e., atomic resolution, but these are limited mainly to static snapshot information without time resolution. On the other hand, time-resolved spectroscopies, fluorescent imaging, and others can directly observe temporal dynamics of proteins, but their spatial resolution is limited. Among others, the high-speed atomic force microscopy (HS-AFM) has a unique feature of directly observing biomolecular structural dynamics near physiological condition at single-molecule level.^1 –7^ Yet, its resolution both in time and space is limited; typically, ∼ 1 nm in lateral direction and ∼ 100 ms in time. Thus, HS-AFM data alone are not enough to model structural dynamics near atomic resolution.

In this respect, the molecular dynamics (MD) simulation can be considered as a complementary approach since it provides very high spatiotemporal information; ∼ 1Å in space and ∼ 1 fs in time. However, MD simulations are limited in reachable timescales; normally, ∼ μs for the standard all-atom model with normal computers.^8,9^ More importantly, MD simulations are based on force fields, which merely approximate to the “true” atomic forces. Any observables calculated from MD simulations would deviate from experimental data to some degree. Thus, of crucial importance is to systematically correct MD simulation data using some experimental data. In fact, much of recent effort is devoted to develop methodologies to achieve this integration.^10–20^

In this paper, we develop a particle filter method to integrate HS-AFM data with MD simulations. By the combination, we aim at getting the best of both data; obtaining high-resolution structural dynamics insights from experimental data. The particle filter method is one of the sequential Bayesian data assimilation approaches.^21–24^ Compared to other data assimilation approaches, such as the linear dynamical system, i.e., the Kalman filter method and the ensemble-Kalman filter method, the particle filter method offers among the most generic, yet computer intensive approach. In the particle filter method, the probability distribution of structures at a time is represented as a finite set of samples, i.e., particles. For all the particles, MD simulations are used to propagate protein structures for a short time period, termed “prediction”, which is followed by the likelihood estimates of the HS-AFM still image at the time, termed “filtering”. Taking into accounts of the likelihoods, we proceed to the next round. Alternative iteration of the prediction and the filtering results in gaining ensemble of structural dynamics that are compatible with the experimental time-series data. Considering the gap between reachable timescales of all-atom MD and the time resolution in HS-AFM, we chose a coarse-grained (CG) molecular model as the simulation tool, by which we can speed up MD simulations by several orders of magnitude.^25,26^

The particle filter method has successfully been applied to non-linear deterministic dynamical systems with relatively small degree of freedom. Compared to many of these cases, application of the particle filter method to HS-AFM data poses major challenges. First, AFM provides surface height data of molecules bound on the stage plane, which is two-dimensional image data. Thus, the filtering is realized in the measurement space of the pixel numbers, unusually large degree of freedom.^24,27,28^ Second, biomolecular dynamics is Brownian and is intrinsically stochastic. How intrinsic stochasticity is reconciled with the particle filter method is an open question. Recently, Matsunaga et al successfully developed a particle filter method to the FRET data of highly stochastic system of a folding protein.^15^ Third, there is order-of-magnitude difference in the timescales of CG-MD simulations (∼ ps) and HS-AFM measurement (∼ 100 ms). Data assimilation in such a case will be a major challenge.

## THEORY AND METHODS

### Particle Filter: General Theory

The particle filter method, also called the sequential Monte Carlo, is a general Bayesian data assimilation approach to integrate experimental time-series data with a simulation model. We begin with a brief introduction of the general theory (see, for example, references 22 and 23 for thorough description). Throughout, we use a discrete time description so that the time takes integer values, *t* = 0, 1, …, T. Let us denote the molecular structure of interest at time *t* as a state space vector ***x***_*t*_. We model the system time propagation as a stochastic simulation model,

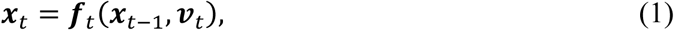

where ***f***_*t*_ defines a system time propagation and ***v***_*t*_ is a system noise vector. Next, we define the experimental measurement at time *t* as a vector ***y***_*t*_, which depends on the state space vector ***x***_*t*_ at the same time and can be written as

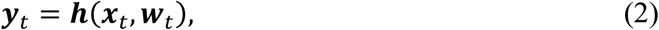

where ***h*** defines the measurement model and ***w***_*t*_ is a measurement noise vector.

In the sequential approach, the molecular system ***x***_*t*_ at time *t* is estimated from the experimental data ***y***_1:*t*_ by the conditional probability distribution *p*(***x***_*t*_|***y***_1:*t*_), where the time-series experimental data from time 1 to t are abbreviated as ***y***_1:*t*_ = {***y***_1,_ ***y***_2_, …, ***y***_*t*_}. Using the Bayes’ theorem, we obtain:

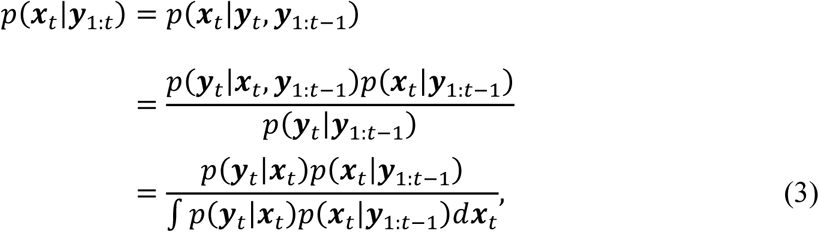

where we used the Eq. (2) from the second to the third line. In this formula, the prior distribution, *p*(***x***_*t*_|***y***_1:*t*−1_), is updated to the posterior distribution, *p*(***x***_*t*_|***y***_1:*t*_), by incorporating the new experimental data, *p*(***y***_*t*_|***x***_*t*_, ***y***_1:*t*−1_), as the likelihood function and the denominator *p*(***y***_*t*_|***y***_1:*t*−1_) as a normalization factor. This procedure is termed “filtering” and *p*(***x***_*t*_|***y***_1:*t*_) is called the filtering distribution. The prior *p*(***x***_*t*_|***y***_1:*t*−1_) in Eq. (3) can be calculated by the time propagation of the system probability distribution from *p*(***x***_*t*−1_|***y***_1:*t*−1_) as

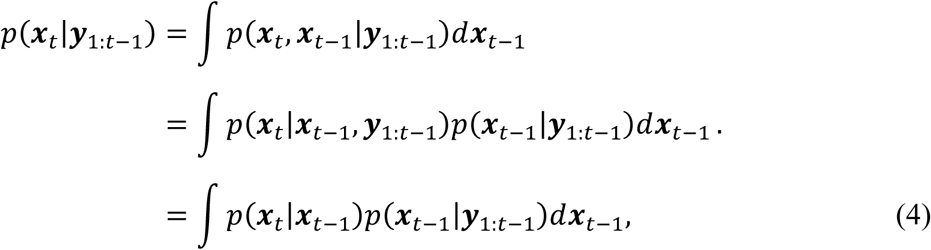

where we assumed ***x***_*t*_ have the Markovian property to reach the third line. Eq. (4) is termed “prediction” and *p*(***x***_*t*_|***y***_1:*t*−1_) is called the predictive distribution. Thus, the time evolution from *p*(***x***_*t*_|***y***_1:*t*_) to *p*(***x***_*t*+1_|***y***_1:*t*+1_) can be performed by one round of the two-step procedure, the prediction followed by the filtering (Figure 1a). By repeating the rounds, we can estimate time series of the distribution sequentially.

**Figure 1:**
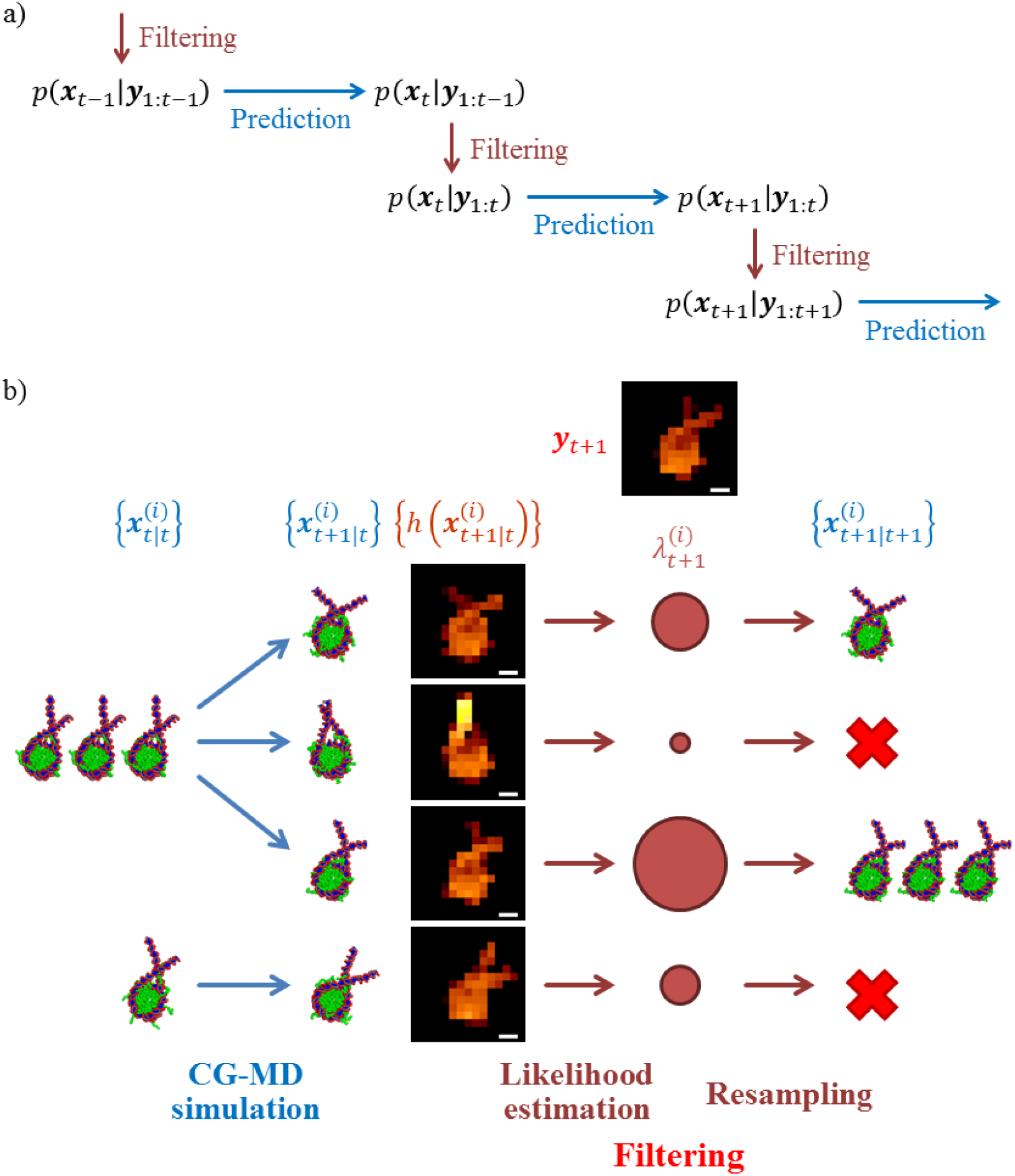
The particle filter method to integrate the HS-AFM data with molecular simulations. a) Sequential update by the two-step procedure, the prediction (horizontal arrows) and the filtering (vertical arrows), in a sequential Bayesian filtering framework. b) Procedure for one round of the particle filter simulation.

In the particle filter method, *p*(***x***_*t*_|***y***_1:*t*_) is approximated by a set of *N* independent samples (called “particles”), 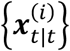, where *i* is a particle index (*i* = 1, …, *N*):

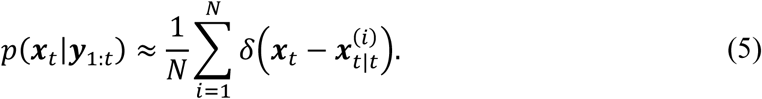

In the prediction step, the simulation for each sample is simply performed and a set of simulation results, denoted as 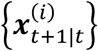, is an approximation to the predictive distribution:

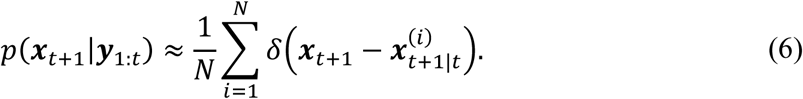

The subsequent filtering step is achieved by estimating the likelihood of each particle *i*, 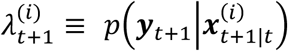, and by drawing *N* particles from the approximate predictive distribution,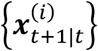, based on the corresponding likelihood, leading to a new set of *N* particles, 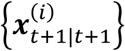, which is an estimate of the filtering distribution, *p*(***x***_*t*+1_|***y***_1:*t*+1_). This process is called “resampling”. Using Eq. (3), *p*(***x***_*t*+1_|***y***_1:*t*+1_) can be approximately expressed as follows:

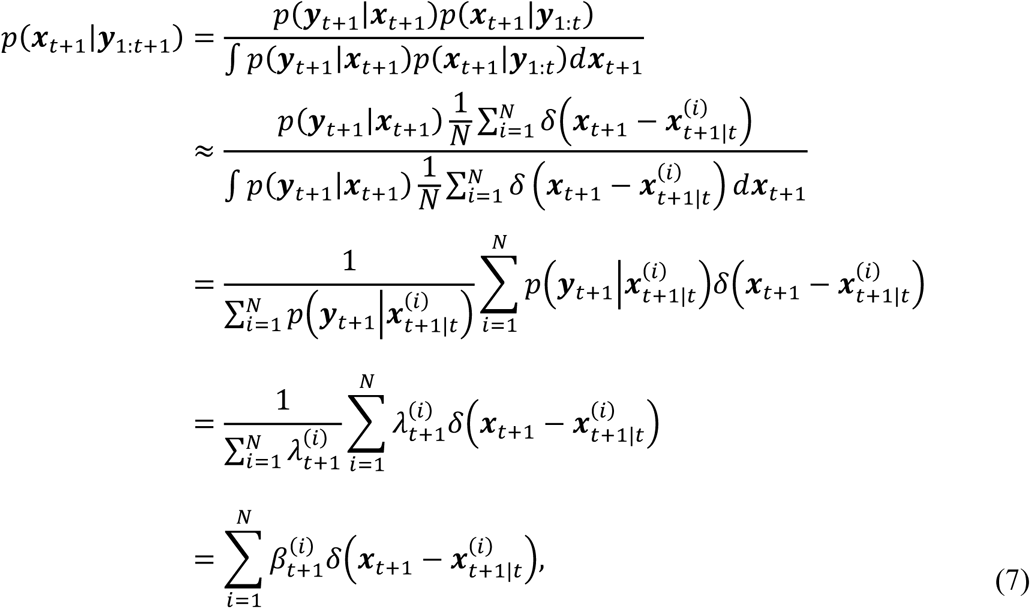

where 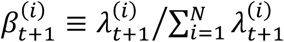, is the weight for the *i*-th particle. By approximating 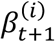 with non-negative integers 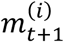 that satisfy the following conditions,

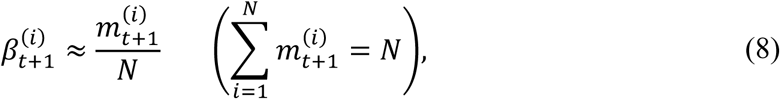

*p*(***x***_*t*+1_|***y***_1:*t*+1_) can be approximated by a set of *N* particles as follows:

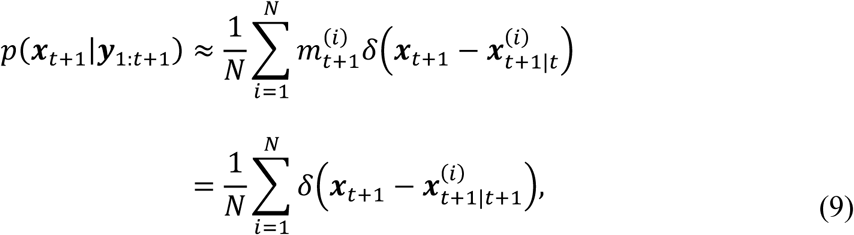

where 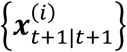 is a set of 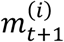-replicated 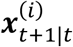:

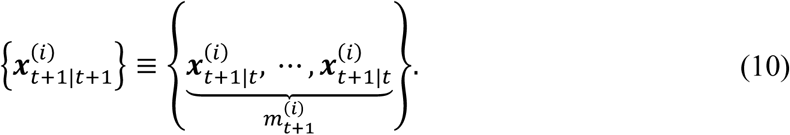

As a result, an estimate of the filtering distribution is likely to contain many copies of a particle with a large weight, while a small-weight particle would not remain in the estimate.

### Particle Filter: Application to AFM Movie with CG-MD Simulation

Now, we apply the general theory to the AFM measurement data that records time propagation of surface shape of biomolecular structures bound on the stage plane. The state vector ***x***_*t*_ collectively represents three-dimensional Cartesian coordinates of all the atoms involved in biomolecular structure (note that an “atom” here means a “CG particle” that represents a group of atoms. Because the term “particle” is used to represent a structure sample in the particle filter method, we avoid to use the term “CG particle”). With our CG-MD method described below, we can simulate the time propagation of ***x***_*t*_. The measurement vector ***y***_*t*_ represents a two-dimensional (the *xy*-plane) AFM image which corresponds to the surface envelope height (the *z*-coordinate) of the biomolecule, stacked in the one-dimensional vector. The dimension of the vector corresponds to the pixel size *N*_*p*_ of the image.

In the present study for HS-AFM data, for simplicity, we assume the linear additive noise ***w***_*t*_ in the measurement model:

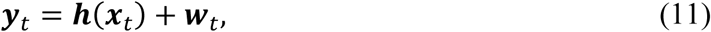

We further assume that ***w***_*t*_ is a white and spatially-uncorrelated Gaussian noise with the standard deviation *σ*. Thus, the likelihood function for the filtering, *λ*_*t*_ ≡ *p*(***y***_*t*_|*x*_*t*_), can be written as

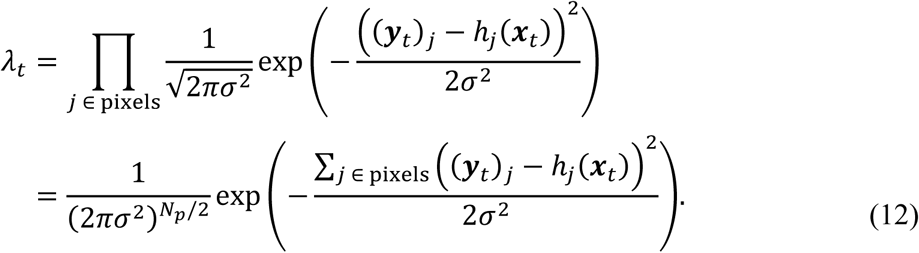

In the particle filter method, the procedure for one round is applied as shown in Figure 1b. A set of *N* particles to approximate the filtering distribution, 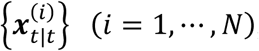, is an ensemble of *N* molecular structures of the target. Using each molecular model as an initial structure, CG-MD simulation is performed for a physical time that corresponds to the discrete time unit. A set of the *N* final structures of CG-MD simulation, 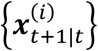, gives an approximated predictive distribution. For the subsequent filtering step, an AFM image of each final structure, 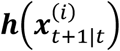, is generated as described below. Then, a likelihood of each particle *i*,

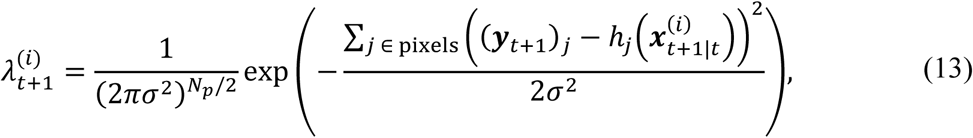

is estimated. Finally, the resampling of particles is carried out by drawing *N* particles with the probabilities proportional to their likelihood, 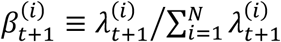, resulting in the next filtering distribution, 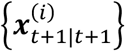. Thus, performing the particle filter simulation by repeating the rounds, we can estimate time series of the approximated distribution of a biomolecule integrating the AFM measurement data.

### Coarse-Grained Molecular Dynamics Simulation

A prerequisite to use the particle filter method is an accurate simulation model for the system of interest. For HS-AFM data, we need a simulation model for biomolecular structural dynamics; in this study, we employ a coarse-grained molecular dynamics (CG-MD) simulation. As a molecule to test the method, we chose a nucleosome, a protein-DNA complex that serves as a fundamental unit of chromatin folding in eukaryotic cells. The nucleosome contains the histone octamer and a duplex DNA of 223 base pairs (bps); 601 strong positioning sequence of 145 bps is flanked by 39-bp linker DNA of the poly-CG sequence in the same way as in our previous work.^29^ The nucleosome is placed on a plane that represents the AFM stage (Figure S1).

We used the CG-MD method that has extensively been used for nucleosomes previously.^29–31^ Since it was described in details, we only summarize it here. Each amino acid in histones is represented by a CG atom located at its Cα position, whereas each nucleotide in DNA is represented by three CG atoms, each representing sugar, phosphate, and base. The histone octamer was modeled by the structure-based AICG2+ potential^32^ and the residue-type-specific excluded volume repulsion. The missing histone tails of histone octamer in the nucleosome were modeled using a statistical potential.^33^ The DNA is modeled by the 3SPN.2C model developed in de Pablo group.^34^ The protein-DNA interactions include the electrostatics in the form of the Debye-Hückel potential, the generic excluded volume interactions, and the sequence-nonspecific hydrogen bond interactions.^29^ Charges in the protein-DNA interaction are defined; −1.0e for phosphate, and acidic residues (Asp, Glu) in the flexible tails, +1.0e for basic residues (Lys, Arg) in the flexible tails, partial charges in the histone octamer core estimated by the RESPAC method.^35^ The all-atom reference structure of the system was prepared by using the crystal structure of the nucleosome (PDB ID: 3LZ0) and an ideal 39-bp segment of DNA.

All CG-MD simulations were performed by CafeMol version 3.2^36^ using Langevin dynamics at a temperature of 300 K with default parameters except for those of excluded volume repulsion, where residue-type-specific radius parameters were uniformly rescaled by a factor of 1.1 to prevent the histone tails from inserting between two DNA strands.^29^ Stage potential of Lennard-Jones type was applied to model the AFM stage and to avoid unrealistic diffusion. The MD step used was 0.3 in the CafeMol time unit. The system was first equilibrated for 10^6^ MD steps so as to stably settle on the AFM stage. We then performed a CG-MD simulation for 10^7^ MD steps with position restraints applied at the centers of mass of three histone multimers, two H2A/H2B dimers and one H3/H4 tetramer, in order not to diffuse on the AFM stage.

To construct AFM images from simulated nucleosome structures, we used the collision detection algorithm implemented in the software, afmize^37,38^ where the probe tip radius is 5 Å and the half apex angle is 5°. When we monitor structural changes of the nucleosome, we used the principal components obtained from the trajectory of CG atoms for phosphate of DNA in the Cartesian coordinates. We note that coordinates of histones are excluded because the histone core structure is always intact, whereas the structure of histone tails is nearly random.

### The Computer Experiment Protocol

We examine the described method that combines HS-AFM data with CG-MD simulation via the particle filter method via a twin experiment.

We first performed a CG-MD simulation for a test system, a nucleosome, at 0.2 M monovalent ion concentration for 10^7^ MD steps, which serve as the ground-truth. In the simulation, the nucleosome lied flat on the stage and its core was sufficiently stable and did not disassemble under the simulation condition. On the other hand, histone tails and linker DNAs were highly flexible and adopted a wide variety of conformations (Figure S2). The histone tails change their conformation very rapidly, but linker DNAs’ movement is so large and slow that it is expected to be observed by HS-AFM. Every 10^6^ MD steps in the simulation, we took the snapshots, from which we constructed a synthetic HS-AFM “measurement” data. The resulting 11 time-point AFM images (including the initial one) constitute the reference AFM movie. To test the effect of the lateral resolution of the AFM measurement, we prepared two sets of reference AFM movies; the resolution of 2 nm × 2 nm with the pixel size 15 × 15 and the resolution of 1 nm × 1 nm with the pixel size 30 × 30.

Then, we performed ten rounds of particle filter simulations using the reference AFM measurement data, with several choices of the number of particles (8, 32, 128, 512, 2048, and 8192), ionic concentrations (0.1 M, 0.2 M, and 0.4 M), simulation time scales, and pixel resolutions of AFM images (2 nm × 2 nm and 1 nm × 1 nm).

## RESULTS

### Particle Filter Simulation to Infer Structural Dynamics of a Biomolecule

To examine the particle filter method for the HS-AFM, we performed a twin experiment. We first created a MD trajectory of nucleosome at 0.2 M salt concentration as the ground truth, from which we made a synthetic AFM movie that serves as the putative measurement data. With this reference AFM movie, we performed the particle filter simulation with 512 particles. The reference AFM movie contains 10 frames, except the initial frame, and thus we alternatively repeat the prediction, Eq. (6), and the filtering, Eq. (7), ten times. Each prediction corresponds to 512 independent CG-MD simulations of 10^6^ MD-steps. Figure 2a plots the trajectories of the 512 particles along the first principal component (PC1) of the ground truth trajectory (blue). During each prediction period, the stochastic nature of MD leads to rapid increase in the structural diversity. In the final structure ensemble of each prediction process, the linker DNAs were oriented in various directions because of their flexibility, while the histone core remained intact. The filtering process begins with the calculation of the likelihood of the reference AFM image for each of structures. The resulting distribution of the likelihood was extremely broad due to the conformational diversity in the linker DNAs (Figure 2b, the constant pre-factors in the log-likelihood is ignored throughout). With these 512 likelihoods as relative probabilities, we then draw a new set of 512 particles, i.e., resampling. We found that, almost always, one particle overwhelmed in the probability (the red trajectory in Figure 2a) so that this single particle occupied the resampled 512 particles, which is often termed degeneracy. We will discuss this issue later more in detail.

**Figure 2:**
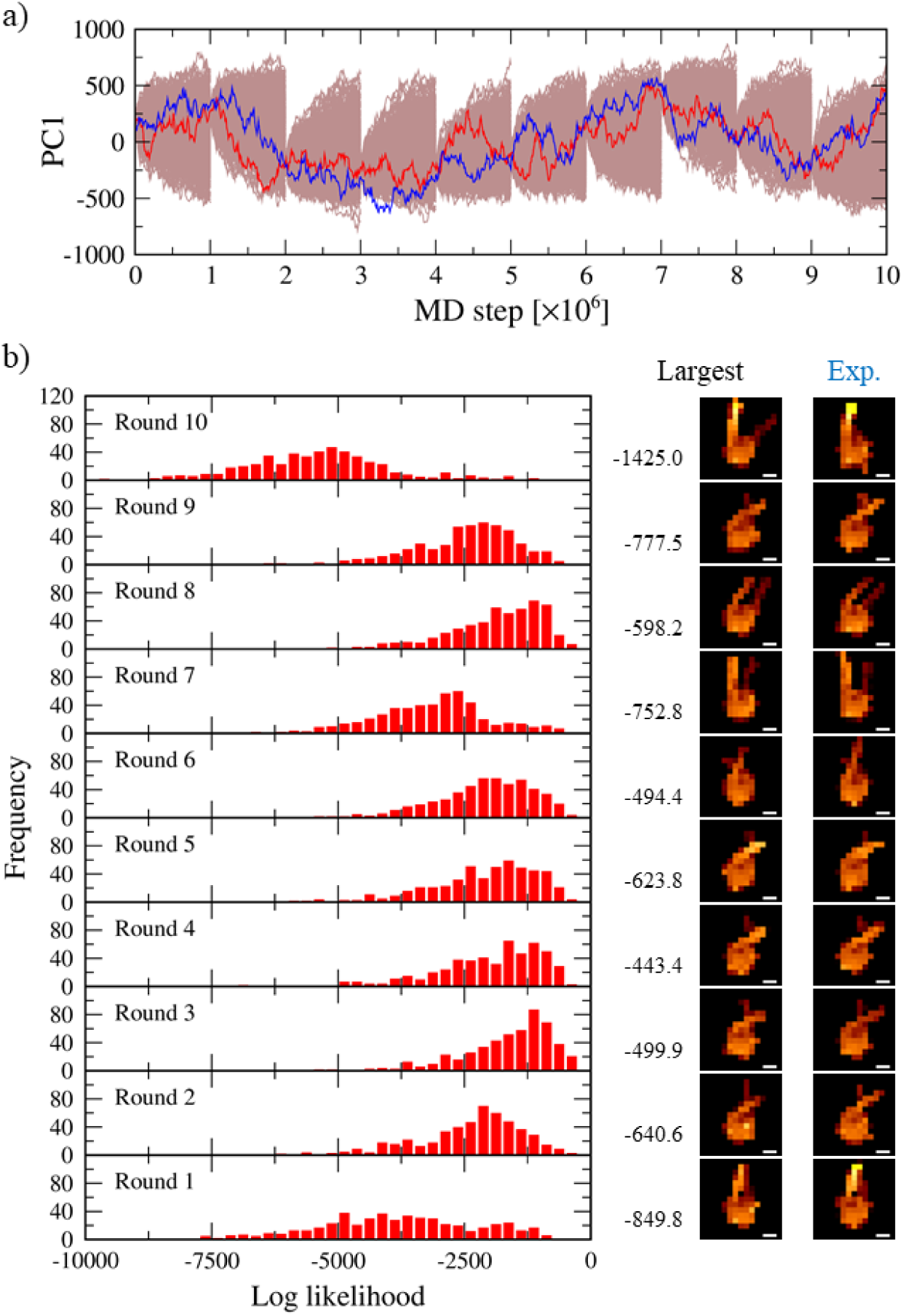
Results of particle filter simulations for a nucleosome with 512 particles. a) Trajectories of the first principal component (PC1) for all particles (brown), the particle that resulted in the largest likelihood (red), and the ground truth (blue). b) (left) Likelihood distribution of particles for each round. The values of the largest likelihood are written on the right side. (right) The AFM images with the largest likelihood for each round (Largest) and of the corresponding synthetic experimental measurements (Exp.) are shown, respectively. Scale bar, 5 nm.

Among the ten rounds, the widths of the likelihood distributions are different (Figure 2b); the distributions in the first and the tenth rounds are much wider than those in the other rounds. This difference seems to depend on the target conformations; the higher the end of the linker DNA is located, the wider the likelihood distribution is. It is probably because the linker DNA moves away from the stage rather rarely and thus this conformation is difficult to be sampled. We confirmed that this tendency of the likelihood distribution is robust by repeating the equivalent particle filter simulations three times (Figure S3). From such a diverse conformational distribution, it is possible to appropriately extract conformations that reproduce the pseudo-experimental results by performing the resampling of the particles based on the estimated likelihoods. In fact, we see that the computed AFM image of the particle with the largest likelihood for each round is similar to the corresponding pseudo-experimental one (Figure 2b right pictures). The values of the largest likelihood for each round show a clear difference, which is understandable due to the sampling difficulty same as the likelihood distribution. Thus, it was confirmed that the developed method can work well as expected.

The total likelihood through ten rounds is simply a product of the likelihood in each round, and the most plausible movement of the target system could be estimated by a series of trajectories with the largest total likelihood (Movie S1). Although, in general particle filter approaches, the whole trajectory that has the largest total likelihood is not necessarily made of the trajectories with the largest likelihood in each rounds, all the three trials in the current work resulted in such a case. The results are quite reasonable because one or few particles with large likelihoods are resampled more frequently from wide likelihood distributions. On the other hand, it is preferable that a certain number of particles are resampled so as to avoid particle degeneracy problems and to well approximate the conformational distribution. In our three trials with 512 particles, however, only a single particle was resampled in almost all of thirty rounds except for some exceptions and this particle degeneracy might be a problem. Various methods have been proposed to solve this problem, but the simplest method is to increase the number of particles.

### The Dependency on the Particle Numbers

Since the accuracy of the particle filter method depends on the number of particles, we now investigate the dependency on the particle number. Especially, the degeneracy problem could be resolved by increasing the particle number. We repeated the otherwise the same particle filter simulations for different numbers of particles; 8, 32, 128, 2048, and 8192, in addition to 512 that is described above.

The results of the most probable trajectory with the largest total likelihood obtained by particle filter simulation with several numbers of particles are summarized in Figure 3. At first sight, comparing the AFM images obtained from the particle filter as the most probable trajectory with those of the reference movie (Figure 3c), we see that those conformations are apparently similar each other in many filtering steps, regardless of the number of particles. Thus, the particle filter method works reasonably in any case. More quantitatively, however, the maximum total likelihood shows the clear dependence on the particle number; the maximum total likelihood increases steadily as the particle number increases (Figure 3a). These results suggest that, not surprisingly, a more accurate result can be obtained merely by increasing the number of particles.

**Figure 3:**
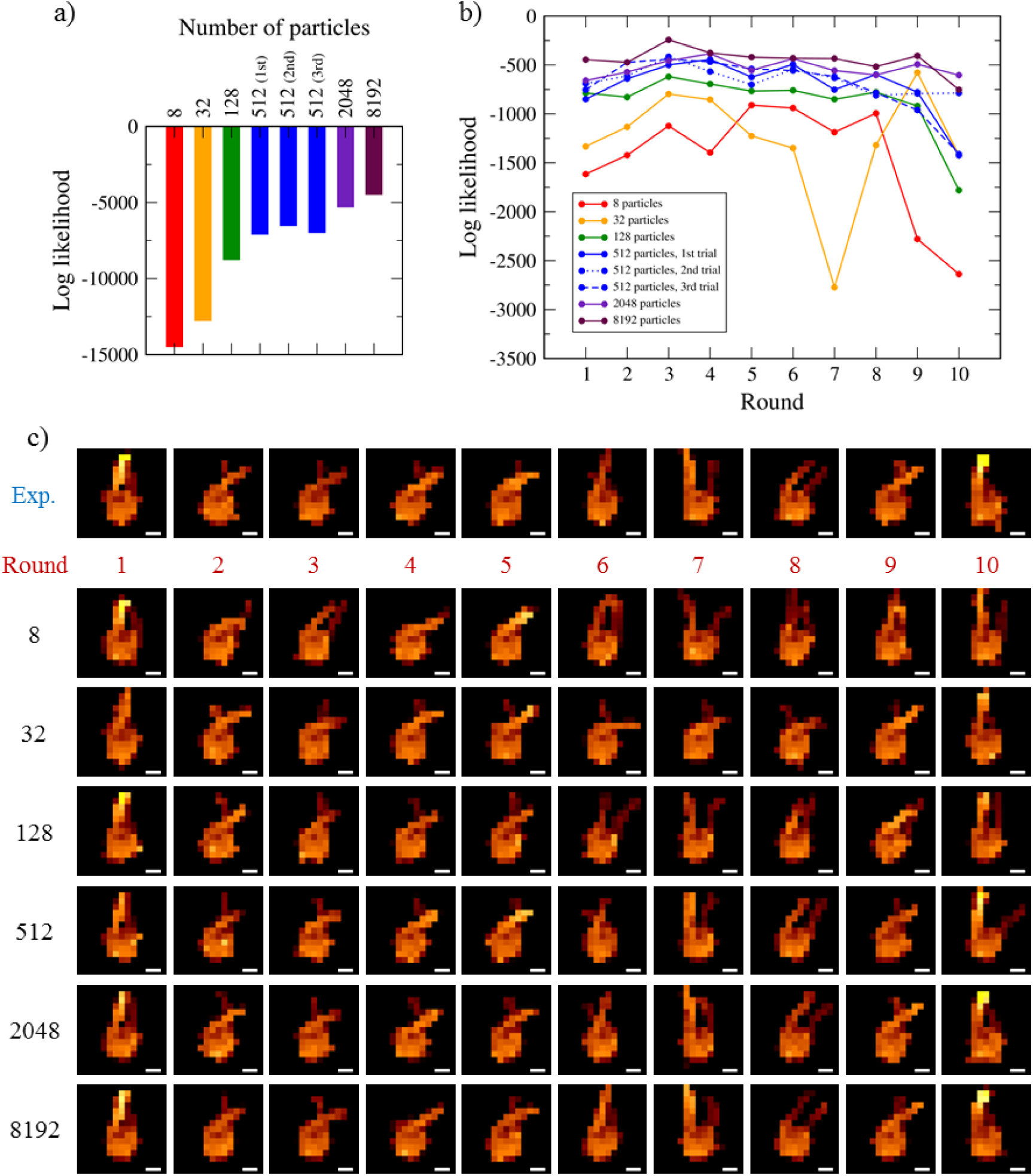
Examination of the particle filter method with different particle numbers; 8, 32, 128, 512, 2048, and 8192. a) The largest logarithm of the likelihood of the whole AFM movie. b) The corresponding likelihood for each round. c) The corresponding AFM image for each round. Only for 512 particles, likelihoods obtained by three trials are indicated separately in a) and b). In c), the reference AFM images (Exp.) are also shown. Scale bar, 5 nm.

Looking into the likelihoods of each round for different number of particles, we found more subtle cases (Figure 3b). In some cases, a fewer particle number led to a larger likelihood, which is opposite to the overall tendency. This reversal of the order seems to be more remarkable for the case of small particle numbers, suggesting that this is caused by large fluctuations associated with small samples. The most prominent example is the comparison between the 8 and 32 particle cases. Up to the fourth round, likelihoods with 32 particles were larger than those with 8 particles as expected, but in the subsequent four rounds, the reversal occurred. In the seventh round, the two linker DNAs in the reference AFM image did not cross each other, which is relatively a rare configuration. With 8 particles, there was a parallel type of structure, by chance, and it was chosen with the largest likelihood particle. But, the structure selected in the case of 32 particles has a crossing of linker DNAs (Figure 3c). As the number of particles is larger, such a reversal is less likely to occur, but it is impossible to prevent it completely. Therefore, when comparing data with different numbers of particles, it is better to discuss results in multiple rounds rather than those in just one round. Alternatively, it is also possible to analyze the distribution of particles in each round instead of one particle with the largest likelihood.

Regarding the degeneracy issue, even with 2048 or 8192 particles, the resampling resulted in only a single particle occupied in all the samples for almost all of twenty rounds. Thus, increasing the number of particles up to 8192 did not resolve the particle degeneracy problem.

### Bayesian Inference of Physical Parameters: Ionic Concentration

One important purpose of the data assimilation, including the particle filter method, is to infer the unobserved (hidden) properties/parameters. In the case of HS-AFM measurement, the behavior of biomolecules can be affected by environmental physical parameters not easy to be observed, such as local ionic strength, interactions to the surface, interaction to the probe tip, and so on. Given the measurement data, one can infer the physical parameter in the framework of the Bayesian approach. Now, we examine whether a physical parameter, the ionic concentration of the solution, can be inferred with the particle filter method for the HS-AFM data. For the purpose, we performed the particle filter simulation of the nucleosome in which CG-MD uses three different salt concentrations, 0.1, 0.2, and 0.4 M monovalent ion concentration. Note that the ground-truth MD trajectory and thus the reference AFM movie were obtained from the 0.2 M ionic concentration. For each of the ionic concentrations, ten-round particle filter simulation with 512 particles was repeated three times.

The MD simulation results showed that the structural dynamics of the nucleosome, especially linker DNAs, was changed according to the ionic concentration. Compared to the case at 0.2 M ionic concentration, at 0.4 M, DNAs in the terminal part of the nucleosome are more unwrapped from the core leading to larger fluctuation. At 0.1 M, on the other hand, the partial unwrapping of the DNAs is reduced, relative to the case of 0.2 M.

The obtained largest total likelihoods through ten rounds depended clearly on ionic strengths as shown in Figure 4a. The largest total likelihood from the 0.2 M case is consistently the largest among the three ionic concentration cases. From these results, 0.2 M can be correctly selected as the most probable value for the ionic strength.

**Figure 4:**
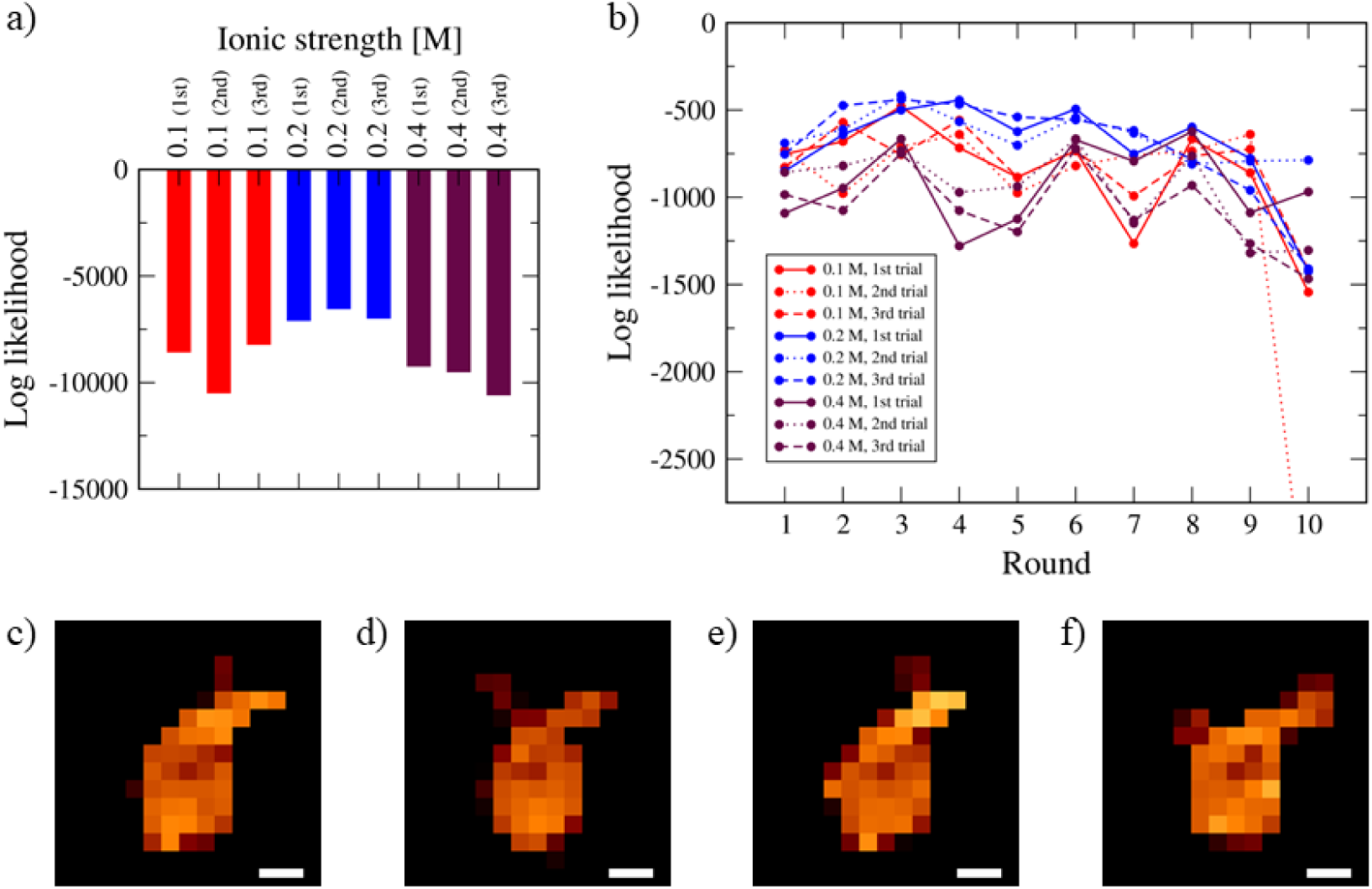
Inferences of ionic concentration via the particle filter simulations. For three trials at each ionic strength (0.1, 0.2, and 0.4 M). a) The largest logarithm of the likelihood of the whole AFM movie. b) The corresponding likelihood for each round. c-f) AFM images in the fifth round of c) the reference and the largest likelihood particle obtained by the first trial at d) 0.1 M, e) 0.2 M, and 0.4 M. Scale bar, 5 nm.

As for the likelihood of each round, however, not all rounds resulted that three largest likelihoods estimated at 0.2 M ionic concentration is larger than those at the other ionic concentrations (Figure 4b). Even in rounds (for example, the fifth round) where likelihood difference is clearly apparent depending on the ionic concentrations, the corresponding AFM images do not seem to have a significant difference (Figure 4c). Thus, the inference of the ionic concentration must be performed by combining the results in multiple rounds.

### Bayesian Inference of Physical Parameters: Timescale

Although the time is a clear physical parameter where there is seemingly no room of inference, we do need to infer the appropriate simulation timescale in the HS-AFM data-driven particle filter approach. First, we note that there is orders-of-magnitude difference in the timescales of HS-AFM measurement and MD simulation. A typical HS-AFM measurement takes ∼10-100 ms per frame, whereas a typical MD simulation uses ∼1-10 fs per MD step. To speed up sampling, we employ CG-MD simulation with a low-friction coefficient and without hydrodynamic interactions. By that, we estimated an effective timescale of a CG-MD step being roughly ∼ 1 ps.^26^ On the other hand, AFM measurements assume modest-strength interactions between the molecule of interest and the stage surface; otherwise, the molecules would move away from the surface by the interaction with the probe tip. These surface-interactions would slow down intrinsic molecular motions to some unknown degree. In CG-MD simulations, we avoided interactions in lateral direction with the surface atoms in order not to slow down the dynamics; otherwise, MD simulations could not simulate large-scale motion comparable to experimentally measured motion. Altogether, when we perform the particle filter simulations for HS-AFM data, the effective simulation timescale relative to the measurement is not a straightforward parameter. Instead, we may need to infer an optimal relative timescale.

Motivated by these arguments, we examine whether it is possible to infer a timescale of pseudo-experimental data appropriately, performing the particle filter simulation on two different timescales, 10^6^ MD steps per round and 10^5^ MD steps per round. We note that the ground-truth MD trajectory contains 10^6^ MD steps per round. Thus, the second case uses an order of magnitude faster timescale than the ground-truth data. For each of the timescales, the ten-round particle filter simulation with 512 particles was repeated three times.

The obtained largest total likelihoods through ten rounds showed an obvious difference depending on timescales (Figure 5a). When the timescale matches, i.e., 10^6^ MD steps per round, all three largest total likelihoods obtained by particle filter simulation are significantly larger than those on the timescale of 10^5^ MD steps per round, as expected. These results suggest that 10^6^ MD steps per round are more suitable than 10^5^ MD steps per round for the time scale.

**Figure 5:**
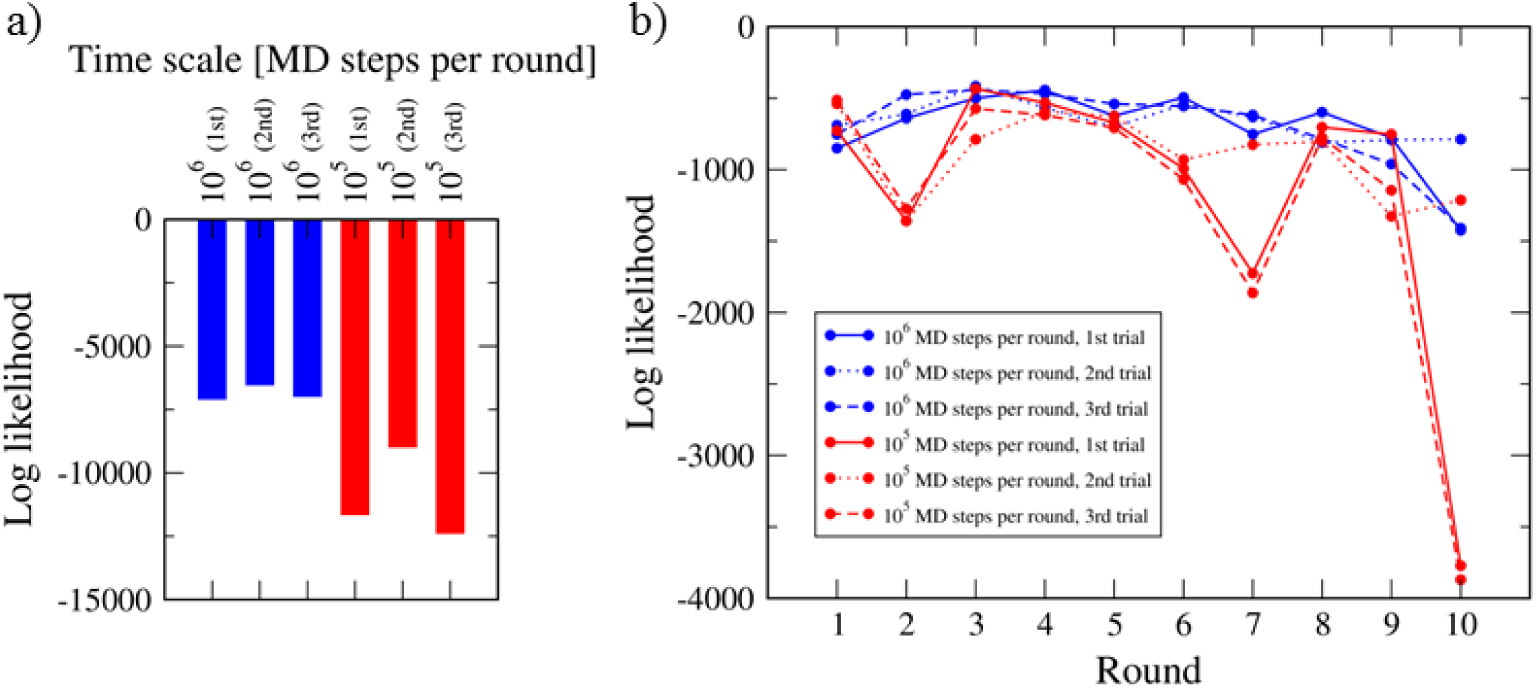
Inferences of timescale via the particle filter simulations. For three trials at each of two timescales, 10^6^ and 10^5^ MD steps per one AFM image. a) The largest logarithm of the likelihood for the whole AFM movie. b) The corresponding likelihood for each round.

On the other hand, the maximum likelihood of each round did not always distinguish the two timescales and can be classified into two types: “distinguishable” and “indistinguishable” (Figure 5b). The second, sixth, seventh, and tenth rounds are of type “distinguishable”, in which there is a significant difference of likelihoods between two timescales, except for seventh and tenth rounds in the second trials on the faster timescale. This difference can be understood that it is difficult to achieve the structural changes observed in these rounds of pseudo-experimental measurement within 10^5^ MD steps. For example, the ground-truth trajectory exhibited marked structural change in DNAs from the first round to the second round (Figure 3c, top line). Such a large change is not easily realized in 10^5^ MD steps. Thus, the maximum likelihood in the second round was markedly small for the case of 10^5^ MD steps per round. In the other rounds, values of likelihood on two timescales are similar and difficult to distinguish (type “indistinguishable”). These results suggest that the target structures can be reached within a fewer number of MD steps than the actual value. Thus, the time scale of the observation should be estimated by using not a single but several rounds. In summary, using ∼10 rounds of measurement, but not in a single measurement, we can robustly infer the correct timescale.

### Effect of Pixel Resolution of AFM Images

The higher pixel resolution of AFM images could provide more detailed information on the structure of biomolecule observed by AFM. Thus, it is expected that the particle filter simulation using AFM images with higher pixel resolution could reproduce biomolecular motion more accurately. Here, we performed particle filter simulation using AFM images with 1 nm × 1 nm pixel resolution, which is twice the original pixel resolution (2 nm × 2 nm), and investigated effects of differences in pixel resolution. For each of the pixel resolutions, the ten-round particle filter simulation with 512 particles was repeated three times.

To our surprise, the obtained largest total likelihoods “per pixel” showed no significant difference between two pixel-resolutions (Figure 6a). Note that the likelihood “per pixel” was used to compare results with different numbers of pixels because the value of likelihood for an AFM image is defined by the product of the value for each pixel as in Eq. (12) and definitely depend on the number of pixels. The likelihood per pixel for each round did not also distinguish two pixel-resolutions (Figure 6b). These results suggest that the use of AFM images with higher pixel resolution do not lead to reproducing more likely motion by the current particle filter simulation. Thus, the original pixel resolution is good enough to distinguish biomolecular structures, leading to estimate their plausible movement successfully. Since this resolution is well attainable in the current HS-AFM experiment, we expect that the current particle filter method works well for actual HS-AFM data.

**Figure 6:**
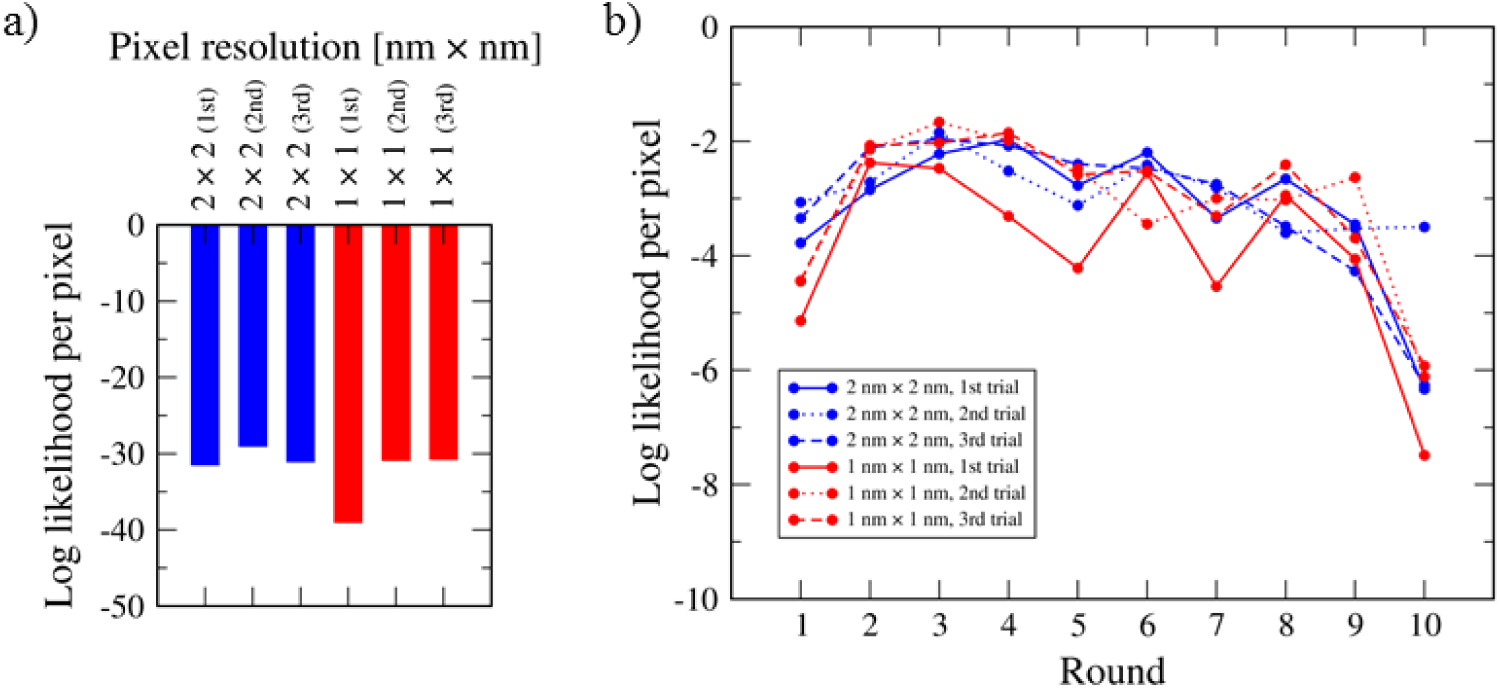
Dependence of the inferences on the pixel resolution of AFM image. For three trials at each pixel resolution (2 nm × 2 nm and 1 nm × 1 nm). a) The largest logarithm of the likelihood of the whole AFM movie. b) The corresponding likelihood for each round.

## DISCUSSIONS

In the current particle filter method for HS-AFM movie, we found that, in the filtering process, the likelihood of one particle overwhelmed all the others and thus the resampling, nearly always, results in the ensemble occupied by copies of the single structure, which is termed degeneracy and is generally regarded as an undesired case in the particle filter method. This severe degeneracy in the current study comes from combination of a few reasons. First, this is in part due to the unusually large degree of freedom in the measurement vector, i.e., the so-called “curse of dimensionality”.^27^ Second, a large gap in the timescales of the HS-AFM measurement of one image and the one CG-MD step: Between the two consecutive AFM images, we need to perform ∼ 10^5^ to 10^8^ MD steps. Third, the intrinsic dynamics of biomolecules is the stochastic Brownian motion: Trajectories deviate as time goes. Altogether, it is obvious that the likelihood of each particle differs in orders of magnitude, resulting in the degeneracy. Even though a straightforward way to avoid the degeneracy may be to increase the number of particles, we found that even the use of 8192 particles does not resolve the issue. For practical use, we cannot increase the number of particles much more.

We note that, albeit undesired, the degeneracy does not directly mean the failure of the particle filter method. In fact, with the given number of particles, we successfully found a high-spatiotemporal trajectory of the nucleosome that is compatible with the reference HS-AFM movie with a certain accuracy. More importantly, our examination of the parameter inference; the ionic concentration and the timescale suggested that we can infer the “true” physical parameter by the likelihood of the entire AFM movie. Thus, we consider that, even with the severe degeneracy, the particle filter method for HS-AFM works well to a certain extent.

However, due to the degeneracy, we need to use the particle filter method for the HS-AFM with caution. For example, in the case of the inference of ionic concentration, we find that, in a single filtering step, the likelihood of that step from the “true” parameter simulation was smaller than that from the wrong one (see the ninth round in Figure 4b, for example). Thus, the parameter inference from a single-step likelihood can be erroneous. Often in other particle filter approaches, people include particles that have different set of physical parameters and, infer these parameter values through the repetitive filtering processes.^39^ When sufficiently diverse particles are picked up in each resampling process, i.e., free from the degeneracy, this way of parameter inference works well as in previous studies. However, with the degeneracy in the current work, this protocol results in the erroneous inference. We need to use multiple rounds of filtering to compare the likelihoods from different parameters. The resampling across different parameters must be performed after sufficiently many rounds of filtering.

A popular alternative to the particle filter method is the ensemble Kalman filter method^40^, which still uses the particle representation in the prediction step, but use the Kalman gain in the filtering process. In general, the ensemble Kalman filter is a powerful method that does not cause the degeneracy issue. Notably, however, in the assimilation to the HS-AFM data, we need a reverse map from the filtered image to a molecular structure. This requires the flexible fitting method of the molecular structure to the given AFM image. Albeit possible^38^, the flexible fitting takes non-negligible computer time, which precludes, in practice, to use the ensemble Kalman filter method in the current problem.

In this paper, we examined the particle filter method for the synthetic HS-AFM movie with promising results, and thus the next step would clearly be its application to real experimental data from HS-AFM. There can be some additional factors need to be considered. First, the real HS-AFM data obviously contain the measurement noise, of which statistical nature is unclear at the moment. The noise would be spatially correlated and the correlation must be un-isotropic; the correlation must be stronger in the x-direction (the direction of the probe scanning) than in the y-direction. This analysis is now underway. Second, interactions between the biomolecules of interest and the stage surface atoms are not homogeneous. Generally, the accurate bottom-up modeling of these interactions may be difficult due to the lack of knowledge of the surface. We may need to infer effective interactions via the Bayesian approach, as was done for the ionic concentration in this study. Third, related to the second point, matching the timescales in the experiment and CG-MD simulations is another factor to be considered (see the first paragraph in the subsection “Bayesian inference of physical parameters: Timescale”).

## CONCLUSION

We developed a particle filter method to combine HS-AFM measurement data with CG-MD simulation. The particle filter method alternatively propagates the structure ensemble by MD simulations of a short time period and then resample the updated ensemble based on the likelihood of the HS-AFM image of that time. With a twin experiment for a test molecule, a nucleosome, we confirmed that the developed method can work well with ∼512 particles. The particle filter simulations could infer the “true” ionic concentration and the “true” timescale by the likelihood of the whole AFM movie.

## ACKNOWLEDGMENTS

We thank Prof. Kazuyuki Nakamura and Suguru Kato for useful discussions. This work was supported mainly by the Japan Science and Technology Agency (JST) grant (JPMJCR1762) (S.T.). The work is also partly supported by Grant-in-Aid for Scientific Research (C) to S.F. (Grant Number 19K06598) from Japan Society for the Promotion of Science (JSPS).

