## Supplemental Figures for "The Particle Filter Method to Integrate High-Speed Atomic Force Microscopy Measurement with Biomolecular Simulations"

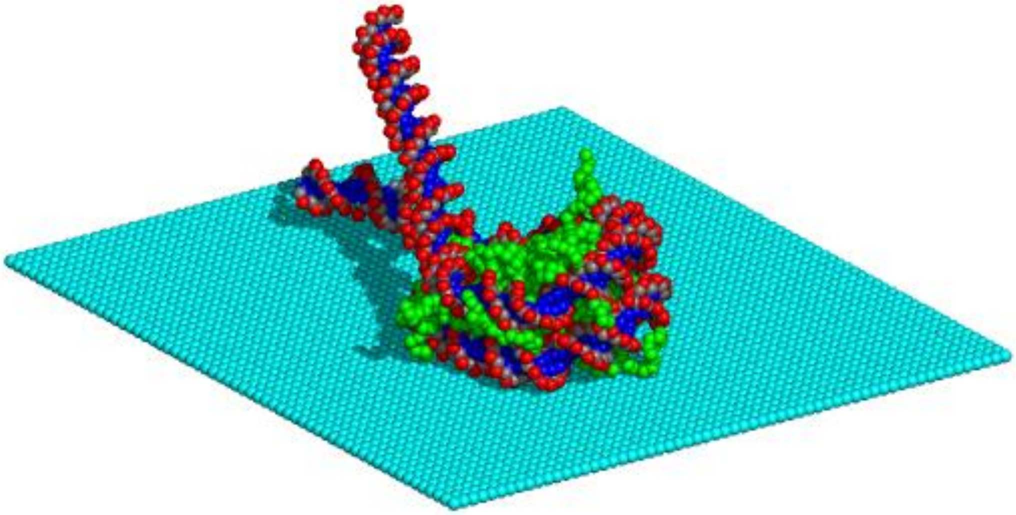

Figure S1: CG model of nucleosome with linker DNAs on the AFM stage.

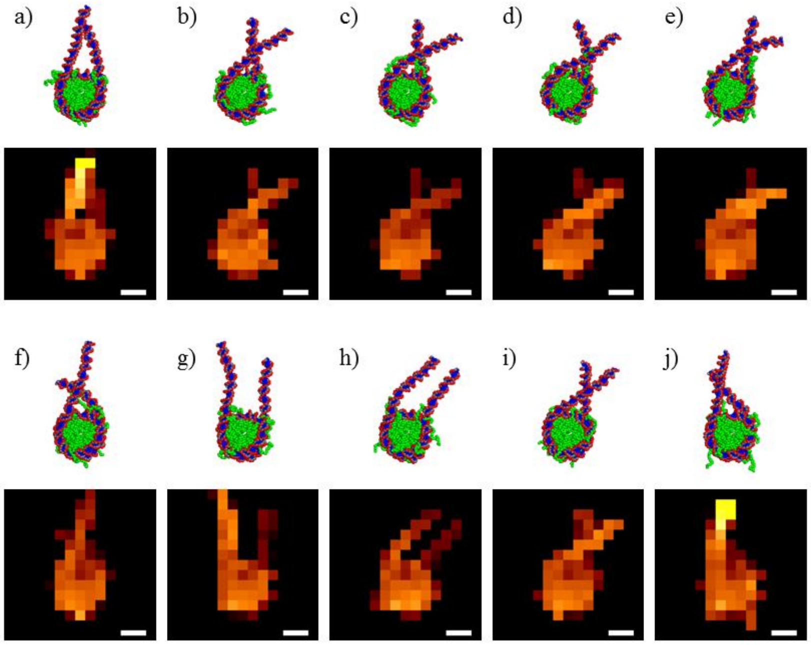


Figure S2: Conformations of nucleosome generated by CG-MD simulation. (a–j) Molecular model (upper) and the corresponding synthetic AFM images (lower) at every $10^6$ MD steps are shown. Scale bar, 5 nm.

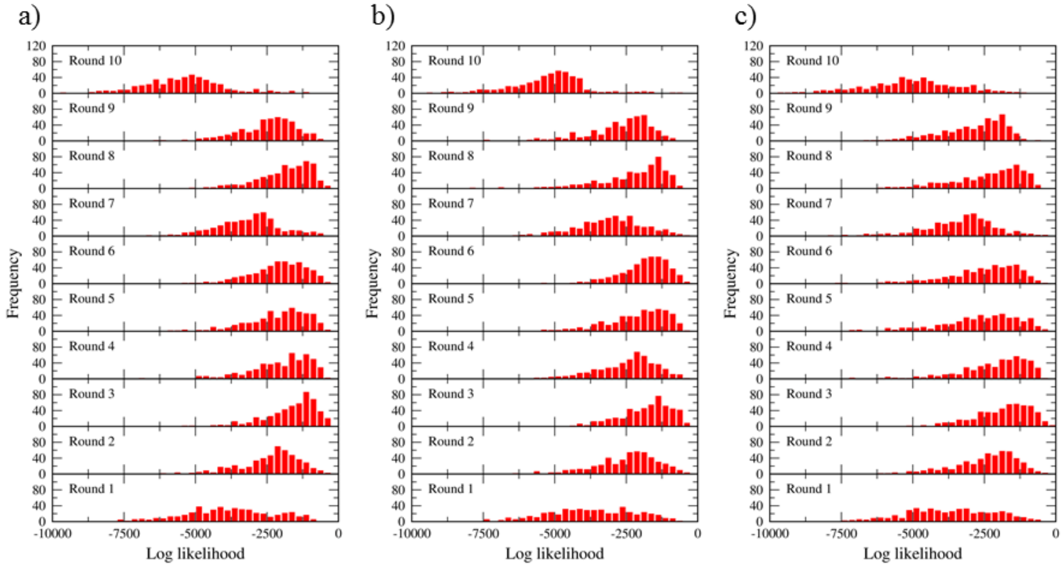


Figure S3: Likelihood distribution of particles for each round obtained by the particle filter simulation with 512 particles. (a) The first trial (same as Figure 2). (b) The second trial. (c) The third trial.

Movie S1: A series of trajectories with the largest total likelihood obtained by the first trial of particle filter simulation with 512 particles. The reference AFM images are shown on the background. The number of MD steps in units of $10^3$ is displayed in the upper left. Scale bar, 5 nm.
